# ExHuMId: A curated resource and analysis of Exposome of Human Milk across India

**DOI:** 10.1101/2020.11.07.372847

**Authors:** Bagavathy Shanmugam Karthikeyan, Janani Ravichandran, S. R. Aparna, Areejit Samal

## Abstract

Human milk is a vital source of nourishment for infants, containing nutrients, immunoprotective components, and bioactive substances. However, several environmental contaminants find their way into human milk. Although lactation physiology has been well documented, the effect of human milk contaminants on maternal and infant health remains unclear. Human milk is the major route of contaminant exposure to infants; these contaminants and their effects can themselves be considered an exposome. While there are chemical regulations in India and scientific literature on environmental contaminants is available, yet there is a lack of systematic compilation, monitoring, and risk management of human milk contaminants. We have harnessed the potential of this large body of literature to develop the <u>Ex</u>posome of <u>Hu</u>man <u>Mi</u>lk across <u>I</u>n<u>d</u>ia (ExHuMId) containing detailed information on 101 environmental contaminants detected in human milk samples, studied across 13 Indian states, compiled from 36 research articles. ExHuMId also compiles the detected concentrations of the contaminants, structural and physicochemical properties, and factors associated with the donor of the sample. Here, we also present findings from a three-pronged analysis of ExHuMId and two other resources on human milk contaminants, with a focus on the Indian scenario. Through a comparative analysis with global chemical regulations and guidelines, we identify human milk contaminants of high concern, such as potential carcinogens, endocrine disruptors and neurotoxins. We then study the physicochemical properties of the contaminants to gain insights on their propensity to transfer into human milk. Further, we employ a systems biology approach to shed light on potential effects of human milk contaminants on maternal and infant health, by identifying contaminant-gene interactions associated with lactation, cytokine signalling and production, and protein-mediated transport. ExHuMId is accessible online at: https://cb.imsc.res.in/exhumid/.

## 1. Introduction

The environmental exposure of women is a concern, especially during pregnancy, and early motherhood (Mead, 2008). A mother is exposed to a myriad of environmental chemicals through food, personal care products, household products, medicines, pollutants, or through her occupational environment (Sonawane, 1995; Landrigan et al., 2002). During motherhood, the practice of breastfeeding is known to be important for the healthy growth and development of a child (Casavale et al., 2019). However, several environmental chemicals, which may affect the child, are capable of entering human milk (Landrigan et al., 2002; Mead, 2008; Lehmann et al., 2018; LaKind et al., 2018). These chemicals are of concern due to the potential impact they can have on maternal health and early development of a child (Leibson et al., 2018; National Research Council (U.S.), 2000). There is a need to monitor, regulate, and consciously avoid these chemicals wherever possible. Biomonitoring of human milk is therefore inevitable (Sonawane, 1995; Landrigan et al., 2002; Mead, 2008; Lehmann et al., 2018; Leibson et al., 2018; LaKind et al., 2018). Environmental contaminants are mostly lipophilic, persistent and bioaccumulative in nature, and have a tendency to deposit in adipose tissues of women or mothers who are exposed to these chemicals (Sonawane, 1995; Mead, 2008). During lactation these chemicals are capable of transfer to human milk majorly via passive diffusion (Agatonovic-Kustrin et al., 2002; Zhao et al., 2006; Heinzow, 2009; Anadón et al., 2011; Vasios et al., 2016). Therefore, human milk is recommended as a core matrix for biomonitoring as per Stockholm Convention on Persistent Organic Pollutants (POPs) (van den Berg et al., 2017) and as a highly efficient non-invasive biospecimen in biomonitoring to detect environmental contaminants (Esteban and Castaño, 2009; van den Berg et al., 2017; Pajewska-Szmyt et al., 2019).

All the environmental exposures of an individual during their lifetime, and their associated health effects, are collectively referred to as the exposome (Vermeulen et al., 2020). In recent times there have been multiple efforts towards the development of large-scale exposome resources. One of the largest exposome databases built to date is the Exposome-Explorer (Neveu et al., 2017) which compiles biomarkers of exposure to dietary and environmental risk factors for diseases. Another manually curated exposome resource, the Human Indoor Exposome Database (Dong et al., 2019), is dedicated to risk factors detected in indoor dust from human exposure studies, and the database compiles 511 chemicals along with their concentrations. T3DB (Wishart et al., 2015) is another exposome database compiling toxic compounds and their target interactions. There has also been efforts toward the development of exposome databases specific to certain biological tissues or biospecimens, such as the Blood Exposome Database (Barupal and Fiehn, 2019).

Given that human milk is a biological matrix, whose monitoring is significant to healthcare and environmental safety, we believe it warrants a dedicated exposome database. The Exposome-Explorer contains a wide range of exposome detected in various biospecimens including blood, urine, plasma, and serum. It also includes the exposures detected in human milk across different geographical regions (Neveu et al., 2017). Some studies have also compiled the list of chemicals detected in human milk, and these studies were published as research articles or scientific reports. A prominent example is the work of Lehmann *et al*. (Lehmann et al., 2018) that has compiled the human milk exposome from samples collected across the United States through literature mining and manual curation.

India is home to a population of nearly 1.33 billion (https://www.statista.com/) with extensive growth in agricultural and industrial sectors, contributing to the production and use of several commercial chemicals in everyday life (Galli et al., 2012). Several studies have detected the presence of environmental contaminants in human milk and a few studies have also compiled the list of chemicals detected in human milk across India (Ramakrishnan et al., 1985; Devanathan et al., 2009, 2012). However, so far there has been no systematic effort towards the monitoring and compilation of these environmental contaminants in India, with the objective to aid chemical risk management and informing policy decisions (Sharma et al., 2014). For example, the report by van den Berg *et al*. (van den Berg et al., 2017) compiles surveys of Polychlorinated dibenzodioxins (PCDDs), Polychlorinated dibenzofurans (PCDFs), Polychlorinated biphenyls (PCBs) and Dichlorodiphenyltrichloroethane (DDTs) in human milk across 52 countries including India. Also a report by Sharma *et al*. (Sharma et al., 2014) compiles the list of some human milk contaminants detected from samples across India. However, these two resources are not comprehensive and the findings are reported at the level of chemical classes, rather than individual contaminants (Sharma et al., 2014; van den Berg et al., 2017). Further, these resources compile only the chemical component of the exposome (Vermeulen et al., 2020), and lack the systematic compilation of maternal factors such as age, body weight, diet, and other factors, which may affect the exposome. This motivated us to develop a systematic approach to compile the <u>Ex</u>posome of <u>Hu</u>man <u>M</u>ilk across <u>I</u>n<u>d</u>ia (ExHuMId), through literature mining and manual curation of research articles that report experimentally detected environmental contaminants in breast milk in studies carried out across India.

We have developed ExHuMId with focus on the Indian scenario. We compile a comprehensive range of information for chemicals detected in human milk samples across India. In addition, we consider a range of additional meta-information, including the geographical location of the sample, and the maternal factors of the donor. All the data in ExHuMId has been standardized, unified, and made accessible through a web interface. ExHuMId is aimed to assist, inform and offer direction to the scientific community, chemical regulators, and the general public, in future research and decision making. ExHuMId is accessible online via: https://cb.imsc.res.in/exhumid.

In this work, we present our comprehensive knowledgebase, ExHuMId, on human milk contaminants in India. We also utilize the information compiled in our resource, and in two other resources (Neveu et al., 2017; Lehmann et al., 2018) to understand the human milk exposome from three perspectives. Firstly, we understand the nature of human milk contaminants through a comparative analysis with several chemical regulatory and guideline lists (Karthikeyan et al., 2020). This has allowed us to identify potential carcinogens, endocrine disrupting chemicals (EDCs), neurotoxins and hazardous chemicals among human milk contaminants. Secondly, we analyze the human milk contaminants based on their physicochemical properties to understand their propensity to be transferred to human milk. Thirdly, we use a systems biology approach (Vermeulen et al., 2020) to predict the effect of human milk contaminants on lactation pathways and cytokine signalling and production. We also investigate possible interactions between the human milk contaminants and xenobiotic transporters which are responsible for the transfer of contaminants into human milk.

We believe that breastfeeding is an essential practice in early childcare, and is one that must be encouraged and made safer (Casavale et al., 2019; van den Berg et al., 2017). Given the ubiquitous nature of environmental contaminants, we believe it is important to be aware and proactive with regards to monitoring, avoiding, and mitigating the risks associated with them. Through the compiled information in ExHuMId and the three-pronged analysis presented here, we aim to shed light on some focus areas that could be taken up in future chemical research and regulatory activity, in order to improve the safety of the much needed practice of breastfeeding.

## 2. Methods

### 2.1. ExHuMId: a curated database of <u>Ex</u>posome of <u>Hu</u>man <u>M</u>ilk across <u>I</u>n<u>d</u>ia

#### 2.1.1. Literature mining and curation

We created the database, <u>E</u>xposome of <u>Hu</u>man <u>M</u>ilk across <u>I</u>n<u>d</u>ia (ExHuMId), with the primary objective of bringing all the published knowledge surrounding human milk contaminants, specific to India, into a single knowledgebase. In other words, ExHuMId compiles the list of human milk contaminants detected in published scientific studies involving samples collected across India. As a first step, we performed an extensive literature search to identify relevant published research articles on PubMed using the following keyword search:

> ((breast OR human OR mother*) AND milk) OR breastmilk) AND India

This keyword search last performed on 24 August 2020, led to 1704 research articles. Subsequently, this set of 1704 articles was manually curated to obtain a subset of articles relevant to the study of human milk contaminants in India (Figure 1). Specifically, we retained only those articles pertaining to “human milk” or “breast milk”, with samples collected only from India. During the manual curation process, we excluded studies on samples collected from outside India, studies without specific geographical indication, review articles or conference abstracts, studies specific to essential nutrients, and articles promoting breastfeeding. This step resulted in a curated set of 36 research articles containing information about the environmental contaminants identified in human milk samples across India, using analytical techniques (Figure 1; Supplementary Table S1).

**Figure 1:**
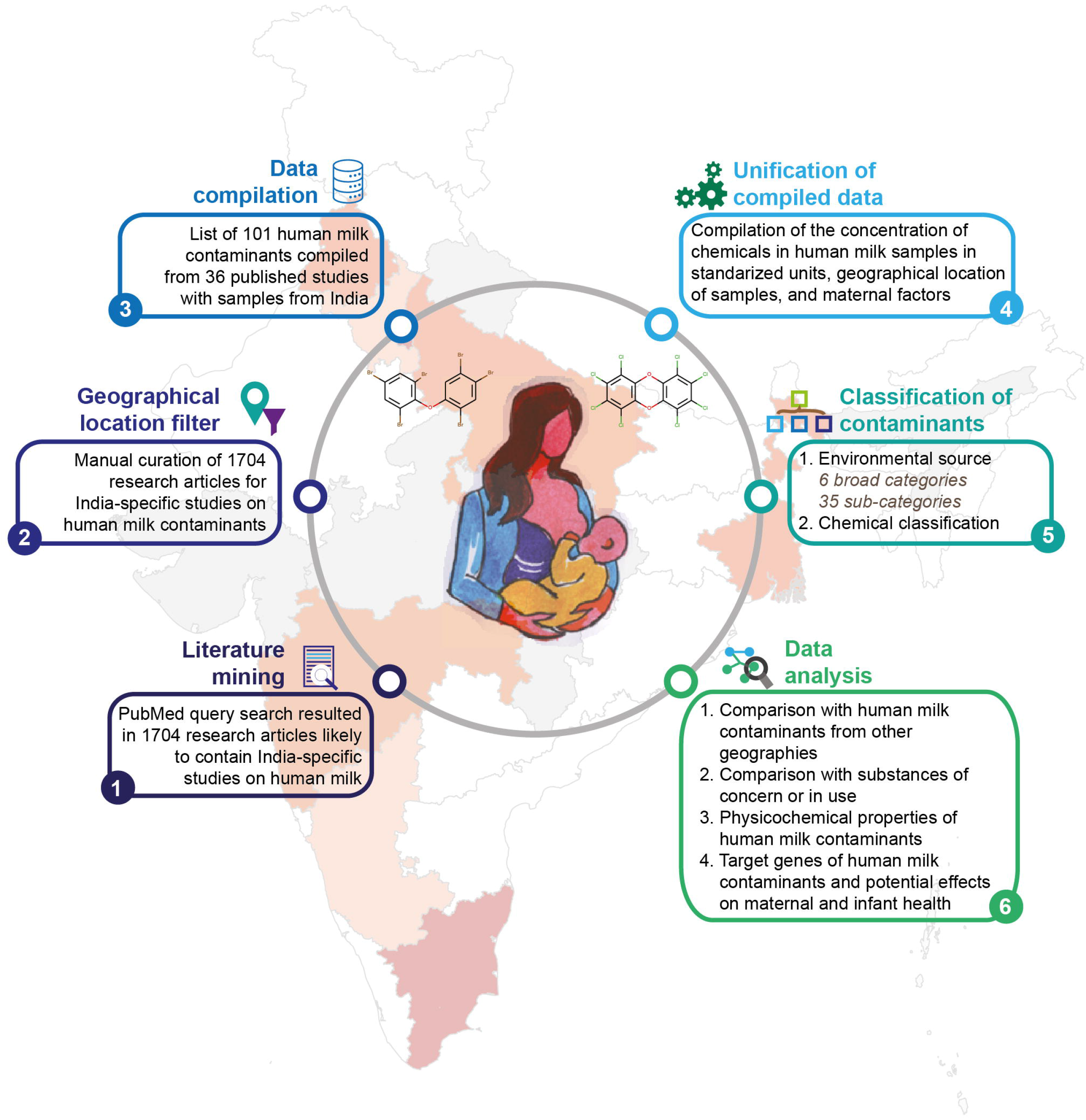
Schematic workflow describing the compilation, curation and analysis of the resource ExHuMId on Exposome of Human Milk across India.

#### 2.1.2. Extraction, compilation, and unification of experimental data

From the curated list of 36 research articles, we have compiled the contaminants including their concentrations detected in human milk samples, geographical location, age, and other factors associated with the mothers from whom the milk samples were collected (Figure 1). For an unambiguous analysis, the data compiled in ExHuMId has been standardized and unified through the following steps.

The first step involved the standardization of the geographical locations from which human milk samples were collected in our curated set of 36 studies. The geographical locations of the study samples were mapped to their respective states in India (Figure 2A). Our manually curated set of 36 studies also recorded a list of maternal factors that influence the presence or transfer of environmental chemicals into mothers’ milk. The second step involved the unification of maternal factors that were compiled from the 36 research articles. The compiled maternal factors observed in the curated set of 36 studies were classified into 9 categories, namely, body weight, food, gestational age, number of pregnancies (Primipara, Biparous and Multipara), occupation, phases of breast milk, residential area, social status, and types of birth (Figure 2B).

**Figure 2:**
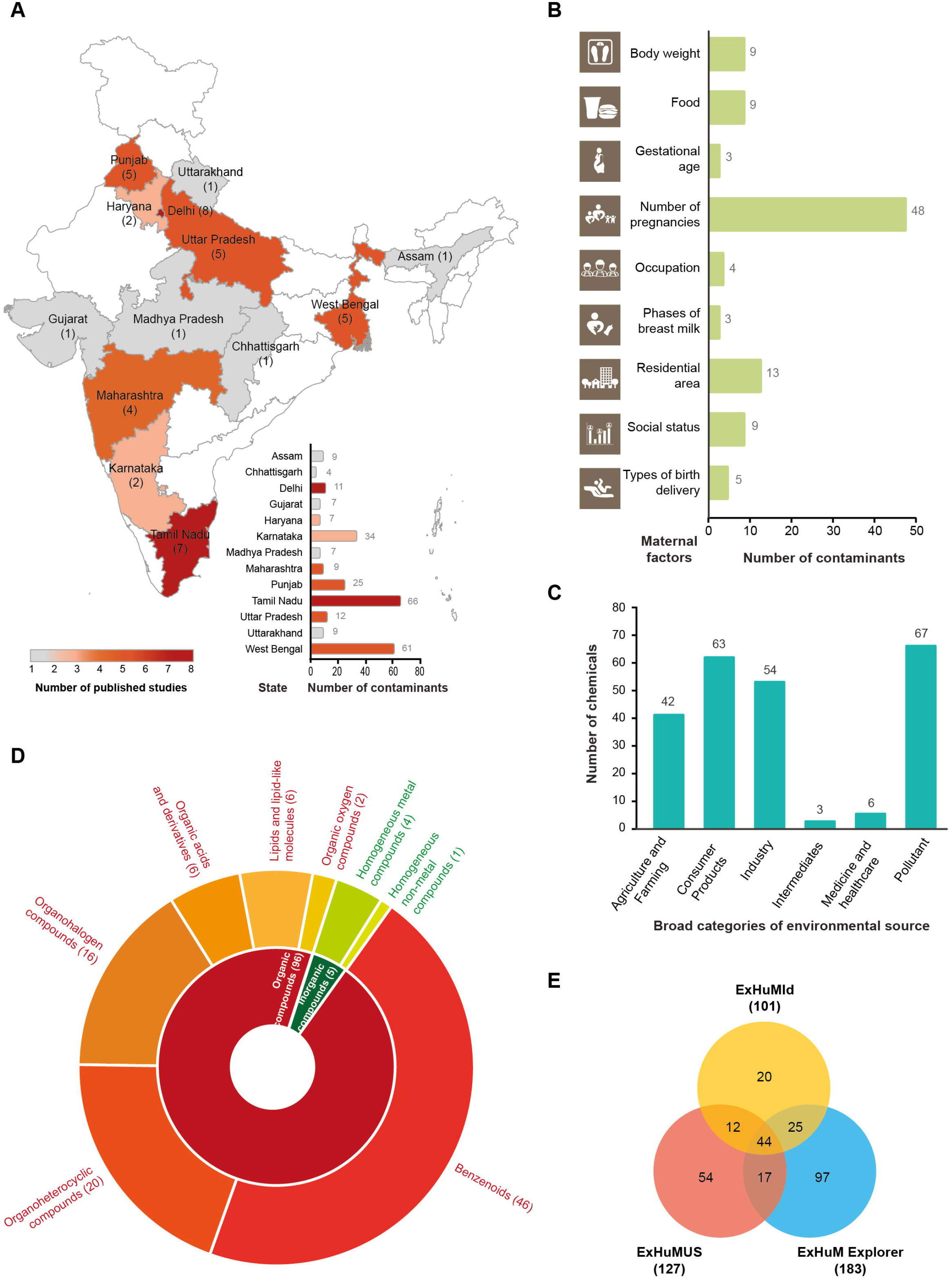
**(A)** An India map displaying different states or geographical locations from where samples were obtained in the curated set of 36 published research articles in ExHuMId on human milk contaminants. The number besides each state in brackets gives the number of published articles reporting human milk samples from that state. The histogram shows the number of contaminants detected across samples obtained from each state. **(B)** Evidence across 9 maternal factors compiled from published articles associated with the human milk contaminants in ExHuMId. **(C)** Distribution of 101 human milk contaminants in ExHuMId across 6 broad categories of environmental sources. **(D)** Sunburst plot showing the chemical classification of 101 human milk contaminants in ExHuMId into 2 kingdoms and 8 super-classes as obtained from ClassyFire. **(E)** Comparison of 101 human milk contaminants in ExHuMId with those in two other resources, namely, ExHuMUS and ExHuM Explorer.

Next, the environmental chemicals detected in human milk across the curated set of 36 studies were mapped to standard chemical identifiers using PubChem (Kim et al., 2019), Chemical Abstract Service (CAS; https://www.cas.org/), ChEMBL (Gaulton et al., 2017), and the Comparative Toxicogenomics Database (CTD) (Davis et al., 2019) to obtain a set of 101 unique chemicals. The final step involved the manual unification of the units for the lowest concentration, highest concentration, mean, standard deviation and standard error associated with the measurement of each chemical in human milk samples in different studies. This step resulted in the unification of the compiled information in 12 different concentration units into 2 standardized concentration units, namely, ug/g lipid weight and ug/L lipid weight.

#### 2.1.3. Classification of human milk contaminants

Following the compilation and standardization of the data on human milk contaminants, we gathered information on their chemical structure including two-dimensional (2D) and threedimensional (3D) structure (in SDF, MOL and MOL2 formats), canonical SMILES, InChI, and InChIKey. We then classified the human milk contaminants based on: (a) their environmental source, and (b) their chemical features. Based on their environmental source, the human milk contaminants have been classified into 6 broad categories (such as, Agriculture and Farming, Consumer Products, and Pollutants) and 35 sub-categories (Figure 2C). The human milk contaminants were structurally classified according to the taxonomy from ClassyFire (Djoumbou Feunang et al., 2016) (http://classyfire.wishartlab.com/), a web-based application (Figure 2D).

#### 2.1.4. Web interface and database management system

ExHuMId is an online knowledgebase that compiles detailed information about the human milk contaminants detected in samples collected from India, with supporting evidence from 36 published scientific studies. This includes their chemical names, unique chemical identifiers, their concentrations as detected in our curated set of experiments, age and maternal factors of the donor of the sample, physicochemical properties, predicted ADMET properties, molecular descriptors, and target genes. Users can also access the identifiers, structural information including 2D and 3D structure for each substance. ExHuMId is accessible at: https://cb.imsc.res.in/exhumid.

The web interface of ExHuMId (Figure 3) was created using PHP (http://php.net/), HTML, CSS, Bootstrap 4, and jQuery (https://jquery.com/). To facilitate interactive visualization, we have used Plotly (https://plotly.com/javascript/), Cytoscape.js (http://js.cytoscape.org/) and JSmol (http://jmol.sourceforge.net/) in the web interface. The compiled ExHuMId database is stored using MariaDB (https://mariadb.org/), and the information from the database is retrieved using Structured Query Language (SQL). ExHuMId web interface is hosted on Apache (https://httpd.apache.org/) web server running on Debian 9.4 Linux Operating System.

**Figure 3:**
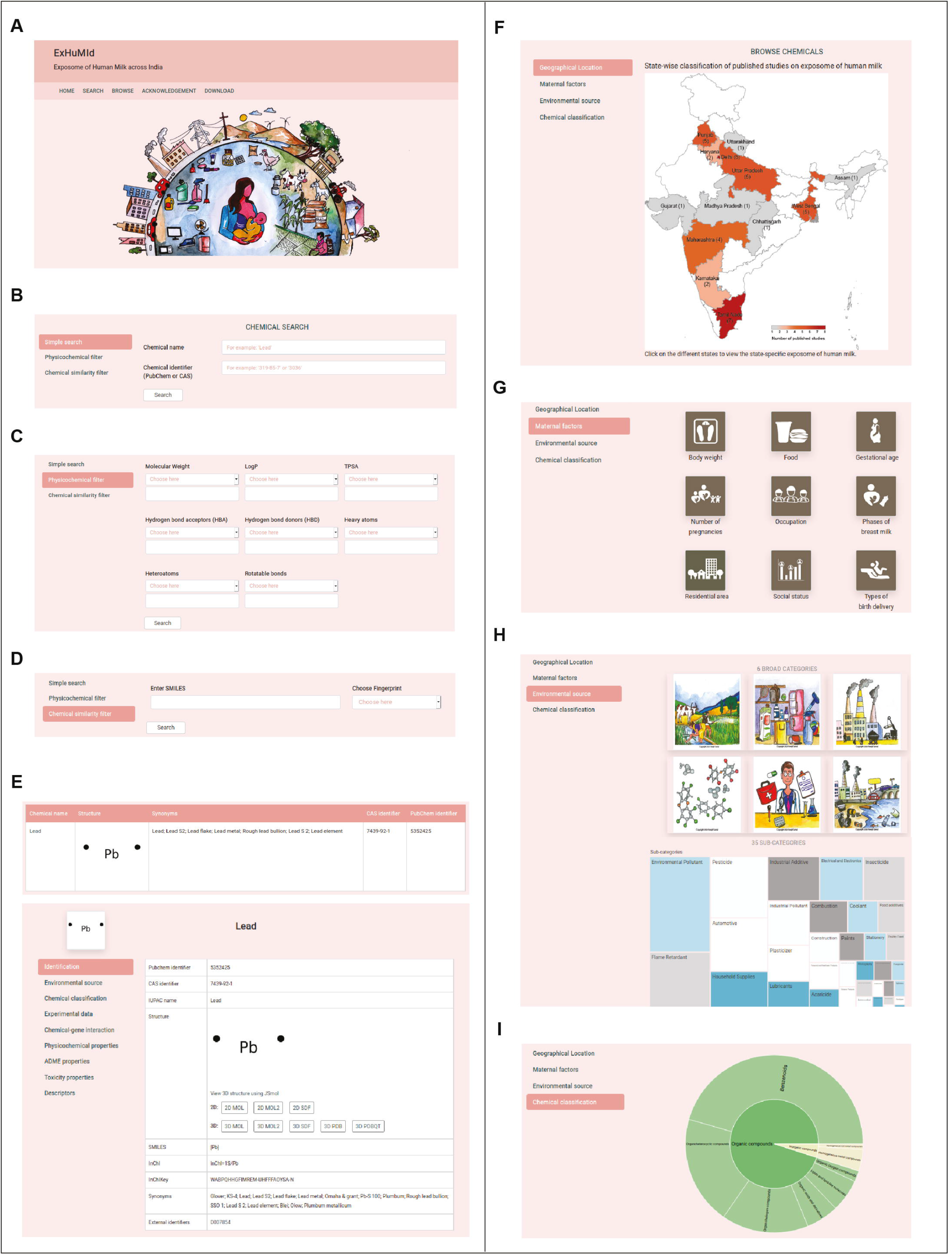
The web interface of ExHuMId. **(A)** A screenshot of the home page of ExHuMId. In Search section, there are three options available to search and obtain information on human milk contaminants compiled in ExHuMId. **(B)** Firstly, Simple search option can be used to search the chemicals using either chemical name or standard identifiers (CAS or PubChem). **(C)** Secondly, Physicochemical filter option can be used to filter the contaminants based on their physicochemical properties such as molecular weight, Log P, TPSA, number of hydrogen bond donors or number of hydrogen bond acceptors. **(D)** Thirdly, Chemical similarity filter can be used to filter the contaminants based on the structural similarity with respect to a query compound. **(E)** The screenshot shows the result page for an individual contaminant. For each contaminant, we can obtain information on structure identifiers, environmental source, chemical classification, experimental evidence, chemical-gene interaction, physicochemical properties, predicted ADMET properties and molecular descriptors. The Browse option in ExHuMId can be used to obtain the human milk contaminants based on: **(F)** Geographical location of samples, **(G)** Maternal factors associated with samples, **(H)** Environmental source classification, and **(I)** Chemical classification.

### 2.2. The Exposome of Human Milk from other geographies

The presence of environmental chemicals in human milk can cause infant exposure to these chemicals, and we here refer to these chemicals as the <u>Ex</u>posome of <u>Hu</u>man <u>M</u>ilk (ExHuM). In order to analyze the environmental chemicals found in human milk, we have considered data from 3 sources. The chemicals in our resource, ‘ExHuMId’ (Exposome of Human Milk across India), have been considered for their specificity to India. The chemicals studied by Lehmann *et al*. (Lehmann et al., 2018) have been considered for their specificity to USA, and we refer to this chemical space as ‘ExHuMUS’ (<u>Ex</u>posome of <u>Hu</u>man <u>M</u>ilk across <u>US</u>A). Several human milk contaminants are also compiled in Exposome-Explorer (Neveu et al., 2017), and these are not specific to any geography, and we refer to this chemical space as ‘ExHuM Explorer’. Notably, there are 127 and 183 chemicals, compiled from 44 and 31 published research articles, in ExHuMUS and ExHuM Explorer, respectively (Table 1; Supplementary Tables S2). Note that the data compiled in ExHuMUS and ExHuM Explorer are not reflective of the entire US and global populations, respectively. However, given that they are the only compilations of human milk contaminants for geographies outside India, we have considered them in this work. The union of the above-mentioned three datasets gives us a list of environmental chemicals detected in human milk samples from various parts of the world, and we refer to this chemical space as ‘Global ExHuM’ (Supplementary Tables S2). The intersection of ExHuMId, ExHuMUS, and ExHuM Explorer (Figure 2E; Supplementary Tables S2), is the set of chemicals that are of potential concern in the Indian, USA and global scenarios, and we refer to this chemical space as the ‘Common ExHuM’ (Figure 2E; Supplementary Tables S2). The analyses described below will enable us to better understand the nature of the chemicals present in the ExHuM.

**Table 1:**
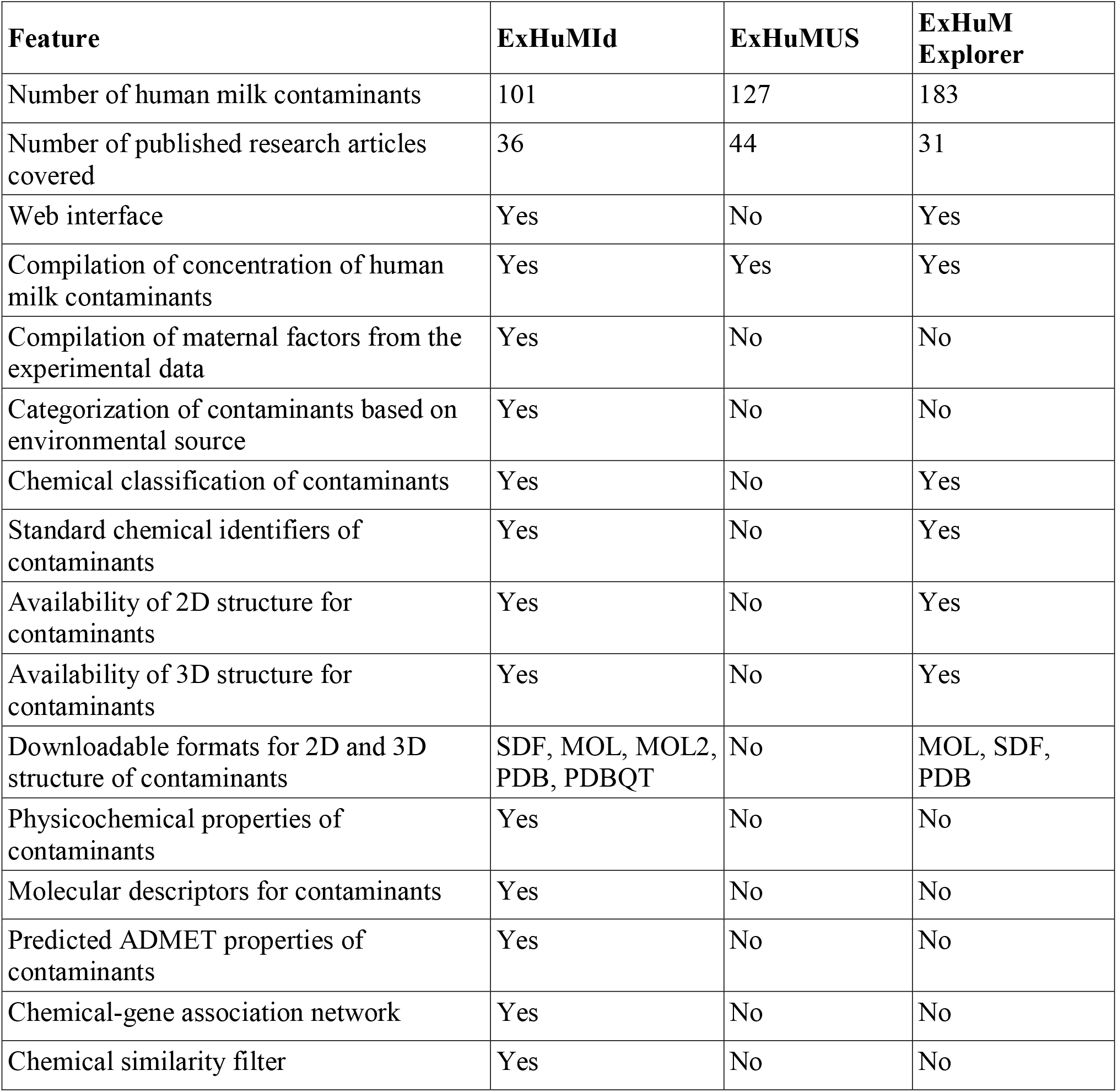
Comparison of the features including meta-information captured in ExHuMId with respect to two other resources, ExHuMUS and ExHuM Explorer, on human milk contaminants.

### 2.3. Analysis of human milk contaminants with substances of concern or in use

We decided to perform a detailed analysis of the chemicals in ExHuM that are of potential concern across the world, with three categories of chemical substances in use or of concern, as described below. The lists of chemical substances employed for this analysis have been described in detail in our recent work (Karthikeyan et al., 2020) (Supplementary Tables S3).

#### 2.3.1. Lists of hazardous substances

These lists consist of known chemical hazards, such as endocrine disruptors, carcinogens, and neurotoxins. Specifically, we have considered four substance lists in this category for analysis of human milk contaminants. Firstly, to understand the presence of endocrine disruptors, we used the list of 792 potential endocrine disrupting chemicals (EDCs) from DEDuCT (Karthikeyan et al., 2020, 2019) for this analysis. Secondly, we considered the list of carcinogens from IARC monographs (Samet et al., 2020). Thirdly, we considered two lists of neurotoxins from the CompTox chemistry dashboard (Williams et al., 2017) of US Environmental Protection Agency (EPA), which are: (a) chemicals demonstrating effects on neurodevelopment (DNTEFFECTS) (Mundy et al., 2015) and (b) chemicals triggering developmental neurotoxicity *in vivo* (DNTINVIVO) (Aschner et al., 2017). Fourthly, we have considered a chemical regulation, namely, the European Union (EU) list of substances prohibited in cosmetic products, and the list was downloaded from: https://eur-lex.europa.eu/legal-content/EN/TXT/?uri=celex:02009R1223-20150416.

#### 2.3.2. Lists of substances specific to India

To obtain a deeper understanding of the contaminants in ExHuMId, we have considered lists that reflect either chemical regulation in India or chemical production scenario in India. Such an analysis is in line with the main focus of this work, that is, Exposome of Human Milk across India. Specifically, we have considered the following lists compiled by relevant departments of Government of India: (a) Production of major chemicals year-wise in India (https://data.gov.in/), (b) List of banned pesticides in India (http://ppqs.gov.in/divisions/cib-rc/registered-products), (c) Schedule 1 hazardous chemicals list in India (http://moef.gov.in/wp-content/uploads/2019/08/SCHEDULE-I.html), and (d) Schedule 3 hazardous chemicals list in India (http://moef.gov.in/wp-content/uploads/2019/08/SCHEDULE-3.html).

#### 2.3.3. Lists of substances causing everyday exposure

A concern and consideration of this study was to better understand the scenario whereby chemicals encountered in everyday life make their way into human milk. For this, we have considered three lists of substances which humans may encounter on a regular basis. We use two lists of substances found in food: (a) FooDB (http://foodb.ca/), and (b) the Joint FAO/WHO Expert Committee on Food Additives (JECFA) list (http://apps.who.int/food-additives-contaminants-jecfa-database/search.aspx). We have also considered two lists of chemicals which are known to be produced in high volume: (a) United States High Production Volume (USHPV) database, and (b) Organisation for Economic Co-operation and Development (OECD) High Production Volume (OECD HPV) list last updated on 2004.

### 2.4. Analysis of physicochemical properties of human milk contaminants

Lipophilic chemicals can be transferred to human milk from maternal plasma via passive diffusion (Agatonovic-Kustrin et al., 2002; Zhao et al., 2006; Heinzow, 2009; Anadón et al., 2011; Vasios et al., 2016). The Milk to Plasma (M/P) concentration ratio is generally used to identify the equilibrium concentration of chemicals in maternal plasma and breast milk (Agatonovic-Kustrin et al., 2002; Anadón et al., 2011; Vasios et al., 2016), and can indicate propensity of the environmental contaminants to enter human milk. However, the M/P ratio, while easily available for drugs, is scarcely available for environmental contaminants (Heinzow, 2009). There is substantial evidence suggesting that the transfer of xenobiotics into human milk is influenced by the physicochemical properties of the chemicals (Agatonovic-Kustrin et al., 2002; Zhao et al., 2006; Heinzow, 2009; Anadón et al., 2011; Vasios et al., 2016). The key physicochemical properties that influence the transfer of environmental chemicals into human milk are the Log P, Topological Polar Surface Area (TPSA), the number of hydrogen bond donors (HBD), the number of hydrogen bond acceptors (HBA), the number of rotatable bonds, and molecular weight (Agatonovic-Kustrin et al., 2002; Heinzow, 2009; Vasios et al., 2016; Zhao et al., 2006). Due to the unavailability of experimentally determined M/P ratio for the 101 chemicals compiled in ExHuMId, we performed a comparative analysis of their physicochemical properties with those of chemicals for which the M/P ratio is available. Specifically, we considered the M/P ratios for a list of 375 chemicals compiled by Vasios *et al*. (Vasios et al., 2016) from published literature, and compared the computed physicochemical properties of chemicals in ExHuM with those compiled by Vasios *et al*. See Figure 4 and Supplementary Table S4 for results from this analysis. The physicochemical properties of the chemicals in ExHuM or Vasios *et al*. were computed using RDKit (https://www.rdkit.org/).

**Figure 4:**
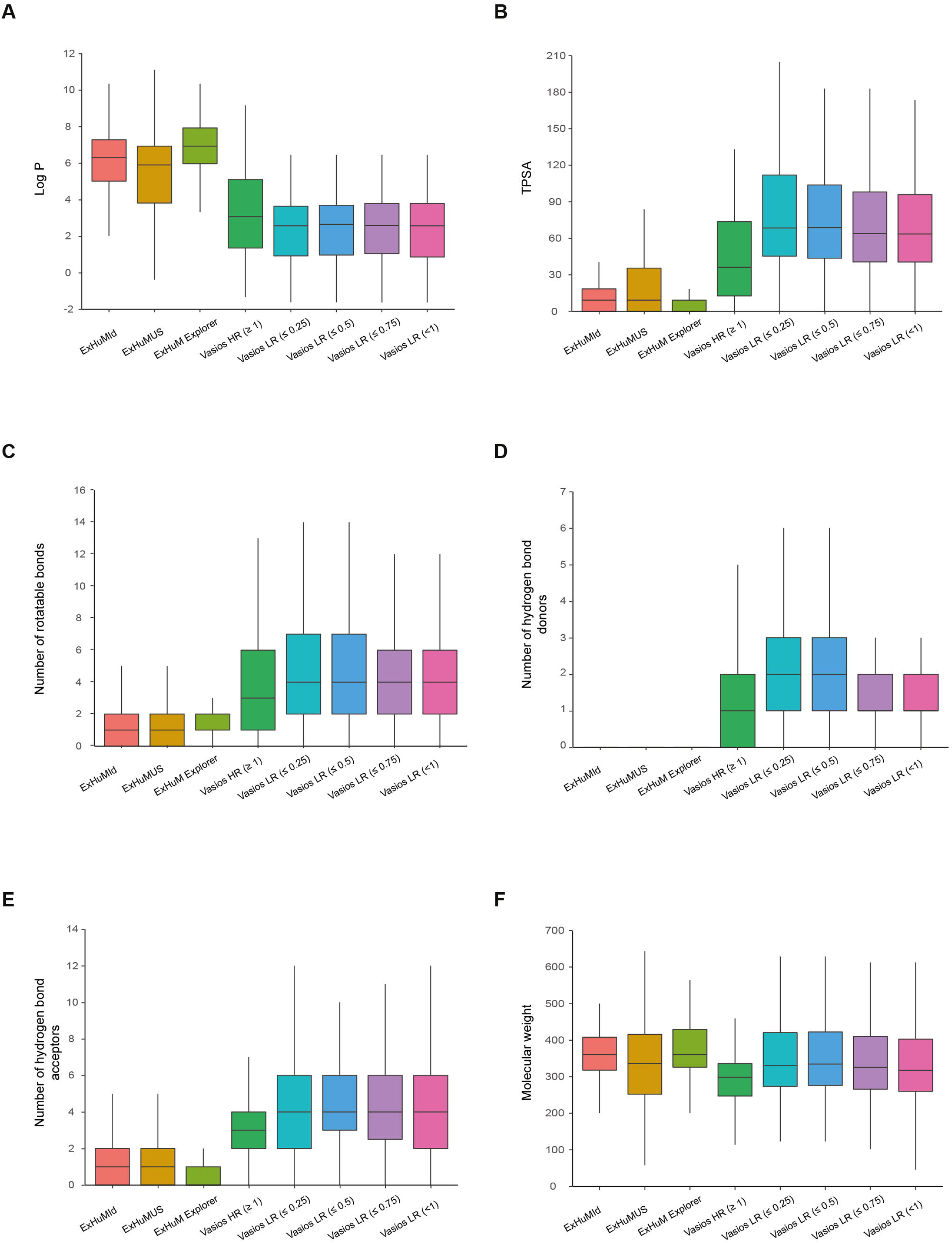
Box plots displaying the distributions of 6 physicochemical properties: **(A)** Log P, **(B)** TPSA, **(C)** number of rotatable bonds, **(D)** number of hydrogen bond donors (HBD), **(E)** number of hydrogen bond acceptors (HBA), and **(F)** molecular weight, for chemicals in 8 different sets, namely, human milk contaminants in ExHuMId, ExHuMUS, ExHuM Explorer, high risk compounds in Vasios *et al*. with M/P ratio ≥ 1 (Vasios HR ≥ 1), and low risk compounds in Vasios *et al*. with M/P ratio < 1 (Vasios LR < 1), M/P ratio ≤ 0.75 (Vasios LR ≤ 0.75), M/P ratio ≤ 0.5 (Vasios LR ≤ 0.5), and M/P ratio ≤ 0.25 (Vasios LR ≤ 0.25). Note that, the distributions for the number of HBD in subfigure **(D)** are not visible for chemicals in ExHuMId, ExHuMUS and ExHuM Explorer as the mean and standard deviation for each of the three sets is very close to 0.

### 2.5. Analysis of potential effects of contaminants on maternal and infant health

Women exposed to environmental contaminants during early stages of pregnancy or lactation have been shown to preferentially store several persistent lipophilic chemicals in their adipose tissue (Mead, 2008), and subsequently during lactation such contaminants can transfer to infants via breastfeeding (Neville and Walsh, 1995; Mead, 2008; Heinzow, 2009; van den Berg et al., 2017; Lehmann et al., 2018; LaKind et al., 2018). In recent times, there have been significant advances in the understanding of lactation physiology and its pathways at a molecular level (Lemay et al., 2013; Maningat et al., 2009) but the effect of environmental contaminants on physiology and health of mother and infant needs further attention (Sonawane, 1995; Landrigan et al., 2002; Mead, 2008). Moreover, the effects of environmental chemicals on adult human beings is better studied, and thus, has more evidence. In contrast, there is a lack of adequate research on the effects of these environmental chemicals on infants, who are exposed to them majorly through breast milk (LaKind et al., 2018; Lehmann et al., 2018). Hence, we were motivated to perform the following analysis to explore the effect of human milk contaminants on mother and child. Specifically, we utilized a network and systems biology approach (Barabási and Oltvai, 2004) to predict the effect of these contaminants on the lactation, cytokine signalling and cytokine production pathways, using existing toxicological resources. In addition, we have investigated if any of human milk contaminants interact with xenobiotic transporters that are responsible for the transport of these chemicals into human milk.

#### 2.5.1. Identifying the target genes of contaminants

To identify the target human genes or proteins of the chemicals in Global ExHuM, we have used two well-known toxicology resources, ToxCast (Dix et al., 2007) and CTD (Davis et al., 2019).

We have used the ToxCast invitroDB3 dataset released in August 2019 (EPA’s National Center For Computational Toxicology, 2019) to retrieve the list of target genes or proteins of human milk contaminants in the Global ExHuM. We used Assay_Summary_190708.csv file which contains the detailed annotation of the ToxCast assays including assay type, assay component, assay component endpoint, assay target information, cell lines used for the assay, and assay citations. For our analysis, we have used the ToxCast assays, and their corresponding assay component endpoints, that are specific to humans. If a tested chemical is found to be ‘active’ for a particular assay component endpoint in a Toxcast assay, then we assign the corresponding gene or protein as the target gene or protein of the chemical.

Thereafter, we also retrieved from CTD the list of target genes or proteins of chemicals in the Global ExHuM using specific filters. In CTD, we have considered only the chemical-gene or chemical-protein interactions specific to humans and those interactions which have at least one evidence in published scientific literature. Moreover, in CTD, we have considered only binary interactions involving one chemical and one gene (Davis et al., 2019), and thus, have filtered out complex interactions. In CTD, we have also not considered the interactions that contained the terms ‘Chemical abundance’ or ‘Response to substance’ based on their ‘interaction actions’.

#### 2.5.2. Identification of contaminants interacting with lactation relevant genes

Environmental chemicals are known to affect the lactation period (Rogan et al., 1987) and the milk secretion (Neville and Walsh, 1995) but the underlying molecular mechanisms by which these contaminants affect lactation physiology and milk secretion remains to be understood. These reported effects on lactation motivated us to investigate if any of the 101 human milk contaminants in ExHuMId can interfere with the genes involved in the pathways associated with lactation.

Prolactin (Hill et al., 1999) and oxytocin (Uvnäs-Moberg and Eriksson, 1996) are the major hormones responsible for lactation. Therefore, we have considered the signalling pathways associated with these hormones for this analysis. We compiled the set of genes involved in the prolactin and oxytocin signalling pathways in humans from NetPath (Chatterjee et al., 2016; Kandasamy et al., 2010; Radhakrishnan et al., 2012) and Kyoto Encyclopedia of Genes and Genomes (KEGG) (Kanehisa, 2000). NetPath compiles a list of genes involved in prolactin and oxytocin signalling pathways in mammals, while the genes retrieved from KEGG are specific to humans. Further, we mapped these genes to their respective human NCBI Entrez identifiers. In this step, we obtained 181 and 237 genes involved in prolactin and oxytocin signalling pathways, respectively, from the above two resources. In addition to these pathways, we have included a set of 14 differentially expressed genes from Lemay *et al*. (Lemay et al., 2013) that are involved in lactose synthesis pathways and important for milk production. Using ToxCast and CTD, we then identified chemicals from ExHuMId that may interact with these lactation relevant genes, as mentioned in the preceding section (Figures 5 and 6; Supplementary Table S6). Moreover, we have also performed the same analysis for chemicals in ExHuMUS and ExHuM Explorer (Supplementary Table S6).

**Figure 5:**
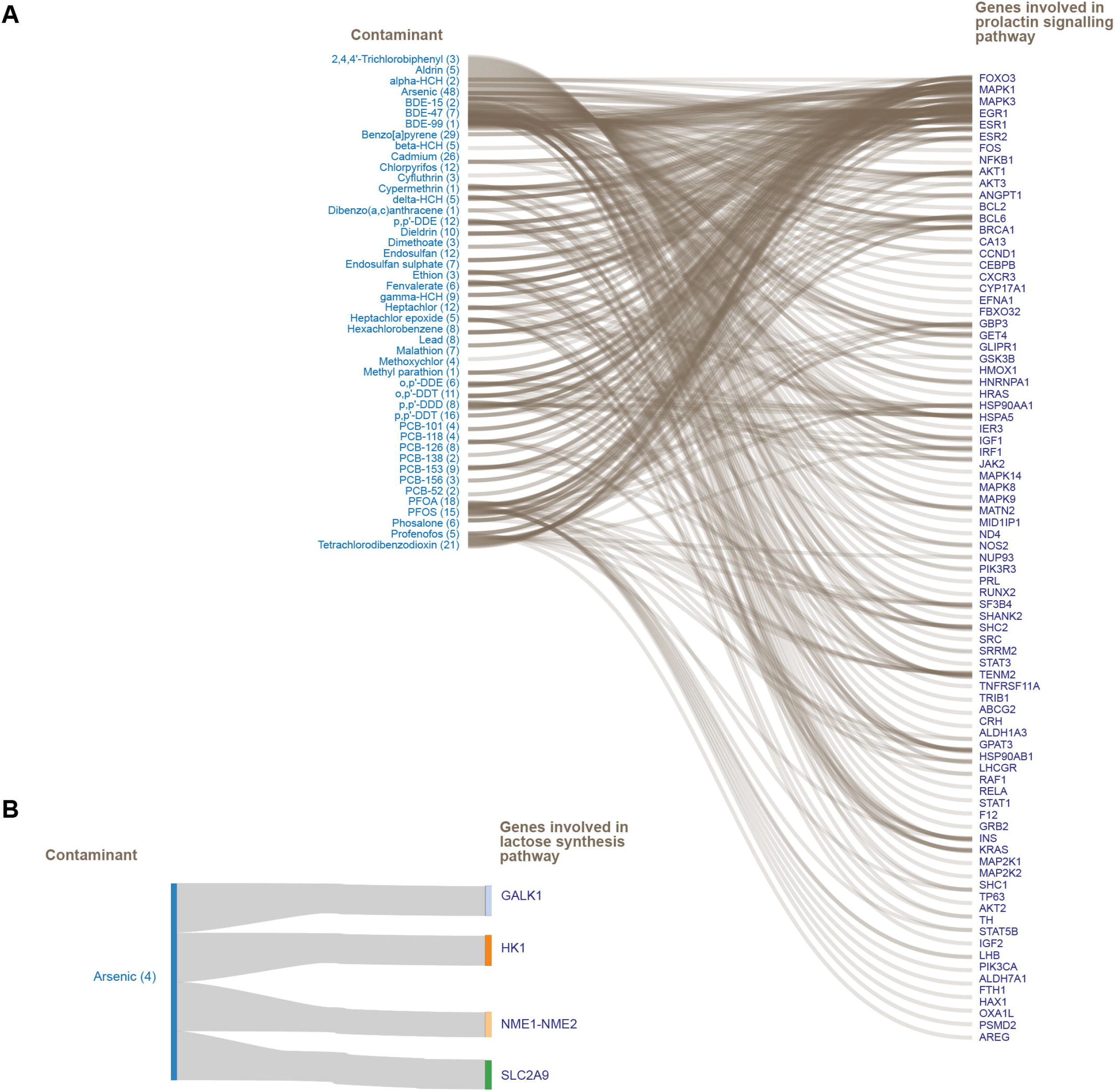
Sankey plots show the human milk contaminants in ExHuMId and their target genes or proteins involved in the pathways affecting lactation: **(A)** Prolactin signalling pathway, and **(B)** Lactose synthesis pathway. Besides each contaminant, the number of target genes is mentioned in parenthesis.

**Figure 6:**
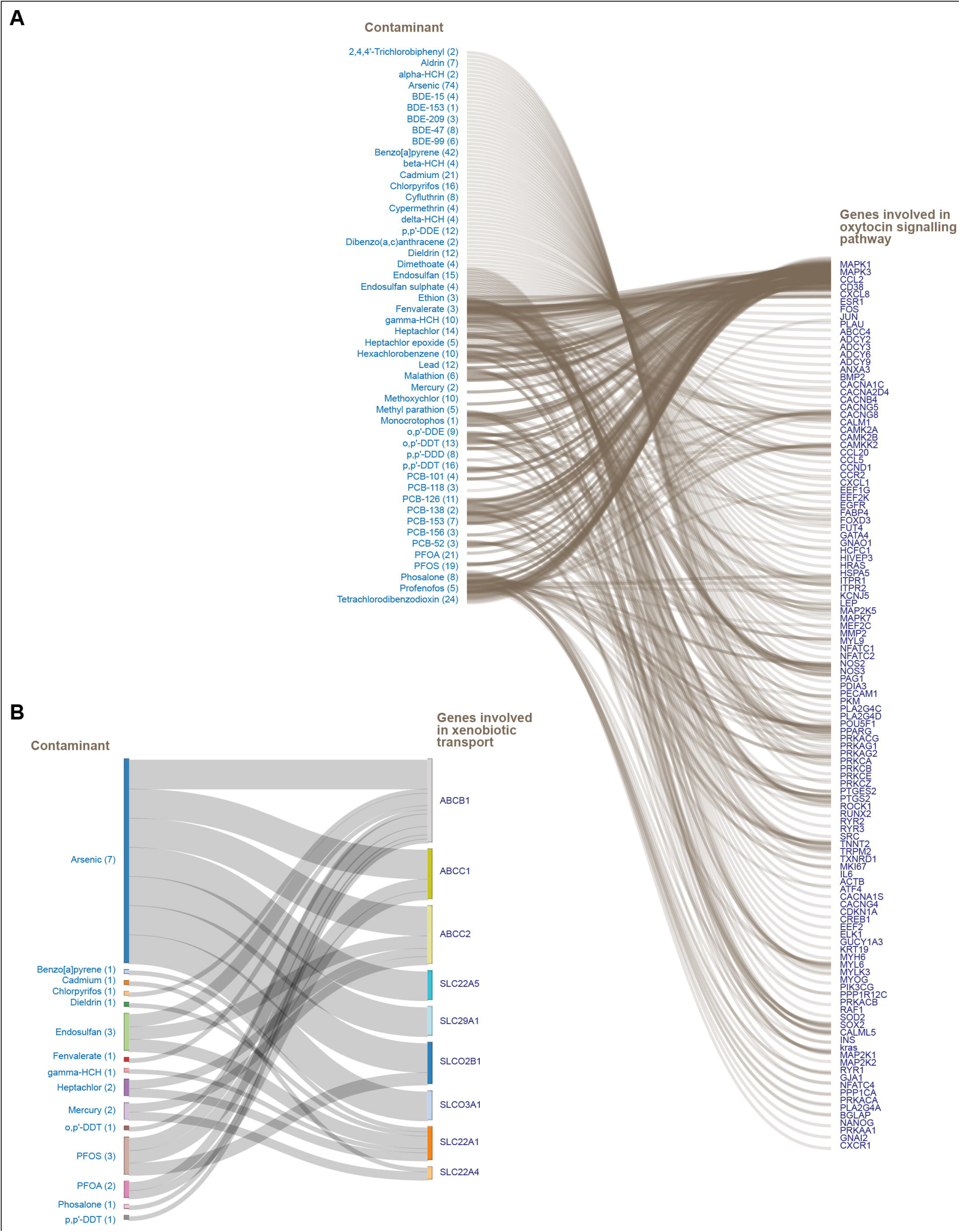
Sankey plots show the human milk contaminants in ExHuMId and their target genes or proteins involved in: **(A)** Oxytocin signalling pathway, and **(B)** Xenbiotic transporters. Besides each contaminant, the number of target genes is mentioned in parenthesis.

#### 2.5.3. Identification of contaminants interacting with cytokine signalling and production relevant genes

Environmental contaminants transferring to human milk were found to be potentially harmful to the development of newborns, due to their ability to disrupt the signalling pathways of infant development (National Research Council (U.S.), 2000; Rebelo and Caldas, 2016; Leibson et al., 2018). Here, we have investigated the effects of human milk contaminants on the immune system development in infants.

It is known that human milk contains several immunological factors including cytokines, chemokines, immunoglobulins, and other soluble receptors that can confer immunity in the lactating infants (Dawod and Marshall, 2019; Jackson and Nazar, 2006). Among these immunological factors, cytokines play a vital role in the regulation of specific and non-specific immune responses (Pajewska-Szmyt et al., 2019). Cytokines bind to the cytokine receptors and trigger the production of cytokines or elicit the immune response via the activation of cytokine signalling pathways (Bagley et al., 1997). Notably, the presence of environmental contaminants in human milk can interfere with cytokine signalling and production (Cohen, 1999; Pajewska-Szmyt et al., 2019). Thus, we aimed to identify chemicals in the Global ExHuM that could potentially disrupt cytokine signalling pathways.

To this end, we first compiled the list of cytokine receptor genes from Cameron *et al*. (Cameron and Kelvin, 2013), HGNC database (Braschi et al., 2019) (www.genenames.org), KEGG BRITE database (Kanehisa, 2000) and Guide to Pharmacology database (Armstrong et al., 2020). In total, we have compiled 116 cytokine receptors for which the chemical-gene interactions were obtained from ToxCast and CTD. Finally, we have gathered the list of cytokines specific to the cytokine receptors that are known to interact with the human milk contaminants. This resulted in a tripartite network containing contaminants or chemicals, cytokine receptors, and cytokines (Figure 7; Supplementary Table S7).

**Figure 7:**
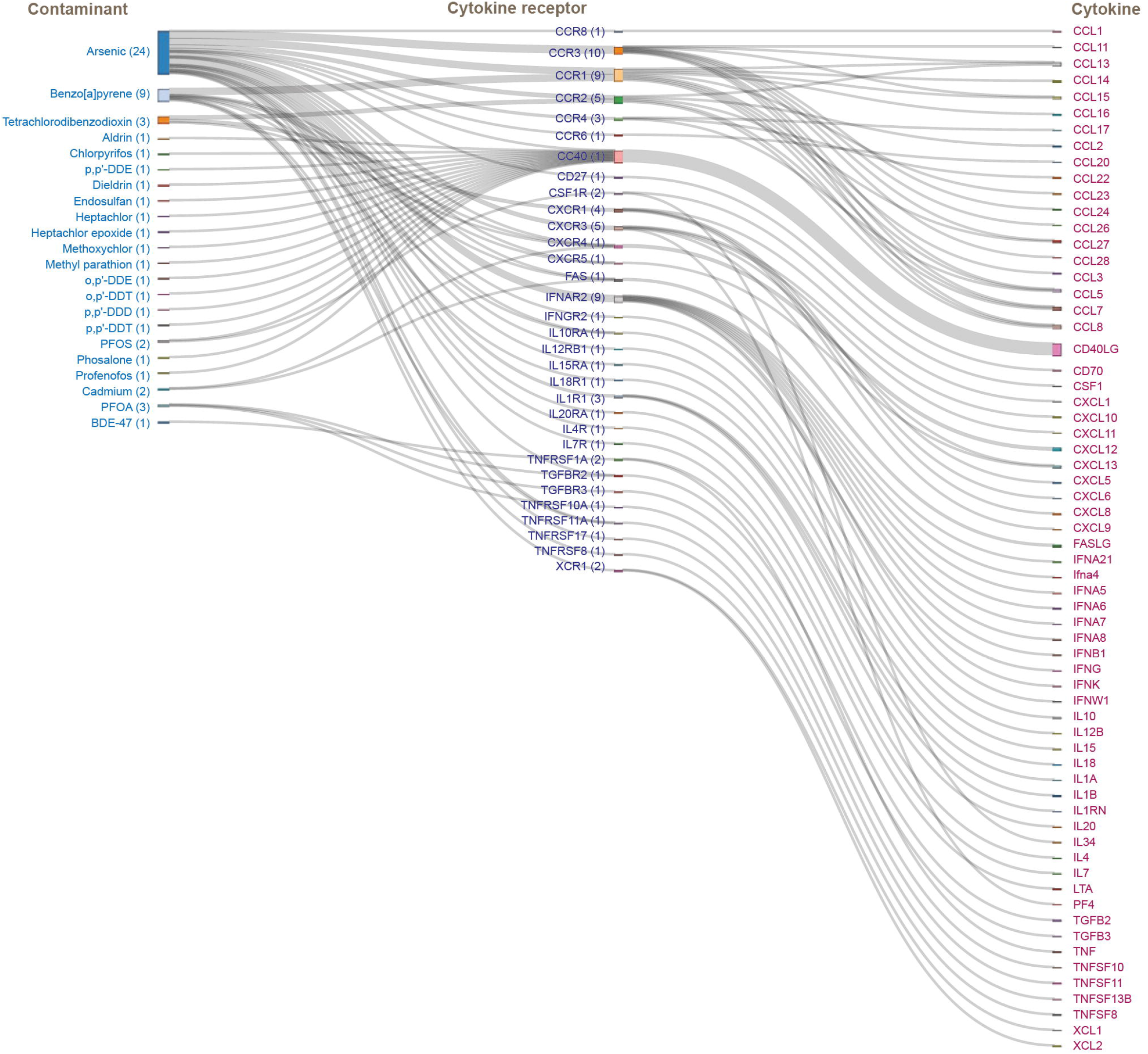
Sankey plot shows the tripartite network of human milk contaminants in ExHuMId, their target genes or proteins corresponding to cytokine receptors, and the cytokines regulated by the specific cytokine receptors. Besides each contaminant, the number of target cytokine receptors is mentioned in parenthesis, and similarly, besides each cytokine receptor, the number of cytokines regulated is mentioned in parenthesis.

#### 2.5.4. Identification of contaminants interacting with xenobiotic transporters

Drug or xenobiotic transporters are membrane proteins that play a major role in transfer of xenobiotics into human milk (Ito and Alcorn, 2003; García-Lino et al., 2019). Some of these transporters have been found to be expressed in mammary gland during lactation (Ito and Alcorn, 2003; Montalbetti et al., 2014; García-Lino et al., 2019; Ventrella et al., 2019). From the study of Alcorn *et al*. (Alcorn et al., 2002), we obtained a list of 19 transporters that are expressed in mammary epithelial cells during lactation. Here, we explore any potential interactions between the chemicals in Global ExHuM and these 19 transporters, using interaction data obtained from ToxCast and CTD (Figure 6; Supplementary Table S8).

## 3. Results and Discussion

### 3.1. ExHuMId: Exposome of Human Milk across India

We believe, in agreement with many others in the science community (Ågerstrand et al., 2017), that scientific knowledge and experimental findings should be readily available to aid and spur further research, inform industry directions and policy decisions, especially when it comes to chemical usage and regulation. Knowledgebases make this possible, by serving as a platform for researchers, industry and regulatory authorities to access a range of useful information. This has motivated us to compile <u>Ex</u>posome of <u>Hu</u>man <u>M</u>ilk across <u>I</u>n<u>d</u>ia (ExHuMId), a curated resource on human milk contaminants specific to India. ExHuMId is accessible online at: https://cb.imsc.res.in/exhumid.

ExHuMId compiles information on 101 environmental chemicals detected in human milk samples studied across 13 states of India, and this information was gathered from 36 published research articles identified via extensive manual curation (Methods; Figure 1; Table 1; Supplementary Tables S1-S2). ExHuMId includes geographical information of the studied sample, age and maternal factors of the donor of the sample, concentrations of the substances detected in the sample, environmental sources of the substance, physicochemical properties, predicted ADMET properties and molecular descriptors (Methods; Figure 2).

Figure 2A shows the distribution of samples collected from each state in India across the curated set of 36 published articles (Supplementary Table S1), and number of chemicals or contaminants detected in each state across the samples. We made an effort to unify a wide range of maternal factors that influenced the presence of contaminants in human milk from the 36 published studies. Among the 9 maternal factors considered here, the number of pregnancies is found to be highly distributed with many more contaminants (Methods; Figure 2B). Note that maternal factors are not available for all samples that have been compiled from the 36 published articles.

Based on the classification of environmental sources, contaminants have been classified into the 6 broad categories ‘Agriculture and Farming’, ‘Consumer Products’, ‘Industry’, ‘Intermediates’, ‘Medicine and Healthcare’, and ‘Pollutant’. The majority of the chemicals compiled in ExHuMId fall under the category ‘Pollutant’ (Methods; Figure 2C). The abovementioned 6 broad categories were further classified into 35 sub-categories based on their environmental sources. Upon classifying the 101 contaminants in ExHuMId based on their chemical class, we find that 96 are organic and 5 are inorganic (Methods; Figure 2D). Among the 96 organic chemicals in ExHuMId, the largest number (46 contaminants) belong to the superclass benzenoids (Figure 2D). We have also unified the concentration units of these human milk contaminants in ExHuMId compiled from 36 published articles. Specifically, ExHuMId compiles 71 chemicals with concentration in standard unit ug/g lipid weight, 18 chemicals with concentration in standard unit ug/L lipid weight, and 11 chemicals with concentrations in both the standard units.

Through the web interface (Figure 3), users can also access the identifiers, structural information including 2D and 3D structure for each human milk contaminant in ExHuMId. The users can navigate ExHuMId via either *simple search* or *browse* options (Figure 3). Using the simple search, users can search for a single chemical in ExHuMId using the chemical name or their PubChem or CAS identifiers (Figure 3B). Users can also search for chemicals in ExHuMId using the *physicochemical filter* or the *chemical similarity filter*. Using the physicochemical filter option in ExHuMId, users can filter contaminants based on their physicochemical properties such as molecular weight, Log P value, TPSA, number of HBA, HBD, heavy atoms, heteroatoms or rotatable bonds (Figure 3C). Using chemical similarity filters, users can search for human milk contaminants similar to a submitted query in standard canonical SMILES format (Figure 3D). Using the Browse section of the web interface of ExHuMId, users can search for chemicals via *geographical location*, *maternal factors*, *environmental sources*, or *chemical classification* (Methods; Figure 3F-I). By navigating to the individual chemical record, users can get detailed information for each chemical compiled in ExHuMId, including the geographical information of the associated sample, maternal factors influencing the presence of the contaminant, concentrations of the contaminant detected in the samples, the structural information and environmental sources (Figure 3E). Users can navigate to the chemical record by clicking the chemical name or its identifiers (Methods).

### 3.2. Comparison of ExHuMId with other resources on Human Milk Exposome

In addition to our curated resource ExHuMId specific to India, we have considered two other compilations of human milk contaminants, namely, ExHuMUS (Lehmann et al., 2018) and ExHuM Explorer (Neveu et al., 2017), here (Methods). Among these 3 compilations of human milk contaminants, we found 44 chemicals to be common, and these 44 contaminants will be henceforth referred to as ‘Common ExHuM’ (Methods; Figure 2E; Supplementary Table S2).

Table 1 presents a detailed comparison of our resource ExHuMId with the other two resources on human milk contaminants. Note that the three resources, ExHuMId, ExHuMUS and ExHuM Explorer, do not have in common any published experimental evidence or literature as the resources compile data on different geographies. Further, the research article (Lehmann et al., 2018) on ExHuMUS provides the list of detected chemicals, their concentrations and the geographical location within USA from where the study samples were collected. However, the ExHuMUS publication is not accompanied by an online resource and the meta-analysis article offers limited information for the compiled list of human milk contaminants (Lehmann et al., 2018). In contrast, ExHuM Explorer (Neveu et al., 2017) contains detailed information on 183 contaminants which were detected in human milk samples collected across several countries. Specifically, ExHuM Explorer gives detailed structural information including the 2D and 3D structures of the contaminants (Neveu et al., 2017). Notably, our resource ExHuMId compiles the different types of information in ExHuMUS and ExHuM Explorer on chemicals, and further, compiles the list of maternal factors that influence the transfer of the contaminants to human milk, their physicochemical properties, their target genes (including visualization of the chemical-gene or chemical-protein interactions), in comparison to the two other resources (Methods; Table 1). In sum, ExHuMId compiles information on human milk contaminants in the specific context of India, and further, makes the compiled information easily accessible to researchers via a user friendly web interface.

### 3.3. Hazardous substances in Human Milk

A better understanding of the nature and exposure sources of the human milk contaminants will most likely help direct further research and regulatory efforts. Here, we focused on identifying substances in the ExHuM that are endocrine disruptors, carcinogens or neurotoxins. These three categories of chemicals are of particular concern due to their potential to affect development and leave behind long-term effects.

#### 3.3.1. EDCs among human milk contaminants

Endocrine disruptors are a major concern due to their potential to interfere with various hormone-mediated processes in the body. Using the list of EDCs in DEDuCT 2.0 (Karthikeyan et al., 2019, 2020), we have analyzed the set of contaminants in ExHuMId and Common ExHuM (Methods). We found that 43 and 19 potential EDCs are present in ExHuMId and Common ExHuM, respectively (Supplementary Table S3). The web interface of ExHuMId provides detailed information on environmental sources of these EDCs detected in human milk samples (Methods).

#### 3.3.2. Carcinogenic substances among human milk contaminants

IARC monographs classify carcinogenic substances into: (a) class 1 that are carcinogenic to humans, (b) class 2A that are probably carcinogenic to humans, (c) class 2B that are possibly carcinogenic to humans, and (d) class 3 that are not classifiable as to its carcinogenicity to humans (Samet et al., 2020). A comparative analysis with the IARC list of carcinogens revealed that 23 and 6 potential carcinogens were in ExHuMId and Common ExHuM, respectively (Methods; Supplementary Table S3). ExHuMId compiles information on 7, 4, 5 and 7 carcinogens belonging to class 1, class 2A, class 2B and class 3, respectively, that were detected in human milk samples from India. IARC also lists a number of commonly found carcinogens, of which 6 were found to be in the Common ExHuM. Thus, 6 commonly found carcinogens having been detected in human milk samples from India, USA, and other parts of the world. Among these 6 commonly found carcinogens in Common ExHuM, there are 3 class 1 carcinogens, namely, 2,3,4,7,8-Pentachlorodibenzofuran, 3,4,5,3’,4’-Pentachlorobiphenyl (PCB-126) and Lindane (Supplementary Table S3).

#### 3.3.3. Neurotoxins among human milk contaminants

Environmental contaminants present in human milk are capable of influencing the neurodevelopment during prenatal and postnatal stages of a child development (Leibson et al., 2018). Based on our comparative analysis with neurotoxins compiled in the CompTox chemistry dashboard (Mundy et al., 2015; Aschner et al., 2017; Williams et al., 2017), we found 14 and 3 potential neurotoxins are present in ExHuMId and Common ExHuM, respectively (Methods; Supplementary Table S3). The 3 potential neurotoxins in Common ExHuM, namely, Chlorpyrifos, Hexachlorobenzene and Lindane are detected in human milk samples collected in India as well as in other parts of the world and are also produced in high volume.

#### 3.3.4. Restricted substances among human milk contaminants

Cosmetic products are a significant source of exposure to various substances, due to their ubiquitous nature and widespread use. The EU chemical regulation prohibits the manufacture and use of a number of cosmetic ingredients within the European Union (Methods). On comparison, we found 16 of these prohibited ingredients are present in ExHuMId, and 4 in Common ExHuM (Methods; Supplementary Table S3). Among these, 3 prohibited cosmetic ingredients, namely, Hexachlorobenzene, Chlorophenothane (DDT) and Lindane are also produced in high volume (Methods; Supplementary Table S3).

EDCs, carcinogenic substances, neurotoxins and prohibited substances have all been identified as hazards, and have been well-studied for their adverse effects. Mitigating the risk posed by these substances will involve identifying their common sources, monitoring and regulating them on a timely basis.

### 3.4. Substances manufactured or regulated in India

We have built ExHuMId with the purpose of compiling and understanding the published data on human milk contaminants from samples specific to India. A comparative analysis of ExHuMId with lists of chemicals manufactured in India and lists from Indian chemical regulations, has further clarified the status of human milk contamination in India (Methods). Several major chemicals manufactured in India have been detected in ExHuMId. Apart from this, 15 substances identified as hazards in Indian chemical regulations are present in ExHuMId, of which 9 are produced in high production volume (Supplementary Table S3). 3 of these 15 major chemicals, namely, Decabromobiphenyl ether, Chlorophenothane (DDT), Lindane, are also present in Common ExHuM (Supplementary Table S3). Further, 9 banned pesticides are also present in ExHuMId. 2 banned pesticides, namely, Chlorophenothane (DDT) and Lindane, are also present in the Common ExHuM, having been detected in human milk samples from USA and other parts of the world (Lehmann et al., 2018; Neveu et al., 2017) (Supplementary Table S3). Further monitoring on the regulatory front and research on the healthcare front may be necessary to mitigate the potential adverse effects of these substances to mother and infants.

### 3.5. Substances contaminating human milk through possible everyday exposure

Humans come into contact with a variety of substances in daily life, particularly via the usage or consumption of an increased number and variety of processed products in today’s world. This is a significant factor in the case of a pregnant woman or breastfeeding mother, since several of these substances may find their way into the mother’s milk (Sonawane, 1995; Landrigan et al., 2002; Mead, 2008). Food additives are one such group of substances to which an average breastfeeding mother is exposed on a daily basis. Especially, the only dietary source of infants is breast milk and presence of any food additives that pose health risk also exposes infants to such risk (LaKind et al., 2018; Lehmann et al., 2018). We found 12 food additives are present in ExHuMId (Methods; Supplementary Table S3), and these include a well-known pesticide Chlorophenothane (DDT). Since Chlorophenothane (DDT) is also present in Common ExHuM, the pesticide has been detected in human milk samples across geographies.

### 3.6. Physicochemical properties of human milk contaminants

Physicochemical properties can influence the transfer of environmental chemicals to human milk from maternal plasma (Agatonovic-Kustrin et al., 2002; Zhao et al., 2006; Heinzow, 2009; Anadón et al., 2011; Vasios et al., 2016), and thus, we decided to analyze ExHuMId from this perspective (Methods). Due to the lack of experimental data on M/P concentration ratios for the 101 contaminants in ExHuMId, we compared the computed physicochemical properties of these chemicals with 375 chemicals with known M/P ratios compiled in Vasios *et al*. (Vasios et al., 2016). Following Vasios *et al*., we have considered the chemicals with M/P ratio ≥ 1.0 as high risk and chemicals with M/P ratio < 1 as low risk for transfer to human milk from maternal plasma. For a more detailed analysis, we have further divided the low risk compounds in Vasios *et al*. based on their M/P ratios into < 1, ≤ 0.75, ≤ 0.5 and ≤ 0.25 resulting in 249, 213, 170 and 114 chemicals, respectively. Thereafter, a comparison of the physicochemical properties was made across the sets of human milk contaminants in ExHuMId, ExHuMUS and ExHuM Explorer, high risk compounds in Vasios *et al*. with M/P ratio ≥ 1, and low risk compounds in Vasios *et al*. with M/P ratio < 1, ≤ 0.75, ≤ 0.5, ≤ 0.25 (Methods; Figure 4; Supplementary Table S4).

Figure 4 shows the mean and standard deviation of the distributions of 6 physicochemical properties, namely, Log P, TPSA, number of rotatable bonds, number of HBD, number of HBA and molecular weight, for chemicals in different sets. In Supplementary Table S5, we report the mean, standard deviation, minimum value and maximum value for the 6 physicochemical properties for the sets of human milk contaminants in ExHuMId, ExHuMUS, ExHuM Explorer, high risk compounds in Vasios *et al*. with M/P ratio ≥ 1, and low risk compounds in Vasios *et al*. with M/P ratio < 1, ≤ 0.75, ≤ 0.5, and ≤ 0.25. We find that the mean and standard deviation of the distributions of 6 physicochemical properties for human milk contaminants in ExHuMId are much closer to those for high risk compounds in Vasios *et al*. with M/P ratio ≥ 1. Note that the high risk compounds in Vasios *et al*. are capable of easily transferring to human milk if they are present in the lactating mother’s body. Further, we observed the same trend for chemicals in ExHuMUS and ExHuM Explorer (Figure 4). Figure 4 also shows a clear difference between the mean and standard deviation of the distributions of the above 6 physicochemical properties for the low risk compounds in Vasios *et al*. in comparison to high risk compounds or human milk contaminants in ExHuMId, ExHuMUS and ExHuM Explorer.

Specifically, the mean lipophilicity (Log P) of human milk contaminants is much higher than chemicals with low risk in Vasios *et al*. For example, the mean Log P of chemicals in ExHuMId is 5.9 ± 2.3 in comparison to 2.4 ± 3.1 for low risk chemicals with M/P ratio < 1 in Vasios *et al*. Moreover, the mean number of HBA, HBD, and rotatable bonds are much lower for human milk contaminants than chemicals with low risk in Vasios *et al*. Also, the mean TPSA of human milk contaminants is much lower than chemicals with low risk in Vasios *et al*. In contrast, there is no clear difference between mean molecular weight for human milk contaminants and chemicals with low risk in Vasios *et al*. In sum, our observations confirm previous observations (Agatonovic-Kustrin et al., 2002; Zhao et al., 2006; Heinzow, 2009; Anadón et al., 2011; Vasios et al., 2016) on physicochemical properties of chemicals with high risk of transfer to human milk from maternal plasma.

Overall, these results give insights into the effect physicochemical properties can have in the transfer of environmental chemicals into human milk, and further, can enable the prediction of such chemicals. While predicting the possible transfer of environmental chemicals into human milk based on physicochemical properties, it is important to bear in mind the due limitations of any such method that does not account for the influence of maternal factors, frequency of exposures, varying pharmacokinetic properties of contaminants, and the complexity of lactation pathways (Sonawane, 1995; Mead, 2008; Lehmann et al., 2018; LaKind et al., 2018; Pajewska-Szmyt et al., 2019).

### 3.7. Contaminant-gene interactions relevant to lactation, cytokine signalling and production, and xenobiotic transport

Though the benefits of breastfeeding outweigh the risk of these environmental chemicals, the effect of these chemicals on mother and infant health remains poorly understood (Sonawane, 1995; Landrigan et al., 2002; Mead, 2008). Using systems biology approach, we provide another perspective from our analysis by predicting the effect of environmental contaminants on lactation, cytokine signalling and production pathways, and xenobiotic transporters with the help of existing large-scale toxicological resources such as ToxCast and CTD (Davis et al., 2019; Dix et al., 2007). Of the 101 human milk contaminants in ExHuMId, information on target genes or proteins is currently available in ToxCast and CTD for 39 and 53 chemicals, respectively. The ExHuMId web interface provides this information on target genes or proteins for different human milk contaminants.

#### 3.7.1. Influence of human milk contaminants on lactation signalling and lactose production

The effect of environmental contaminants on the lactation period (Rogan et al., 1987) and milk secretion (Neville and Walsh, 1995) is well documented. Based on this fact we tried to check if any of the contaminants compiled in the Global ExHuM are capable of interacting with genes involved in lactation signalling and lactose production pathways, by analyzing the prolactin and oxytocin pathway genes (Methods). We found 46 human milk contaminants compiled in ExHuMId target 83 genes out of 181 genes associated with the prolactin signalling pathway (Figure 5A; Supplementary Table S6). In the case of oxytocin signalling pathway, 118 out of 237 pathway-associated genes, are found to be the targets of 50 human milk contaminants compiled in ExHuMId (Figure 6A; Supplementary Table S6). Arsenic targets 48 genes of prolactin signalling pathway and 74 genes of oxytocin signalling pathway. The ESR1 (Estrogen Receptor 1), which is associated with both the oxytocin and prolactin signalling pathways, appears to be a common target, having interactions with the highest number of human milk contaminants (Figures 5A and 6A; Supplementary Table S6). Through the analysis of the genes responsible for the production of lactose, as reported by Lemay *et al*. (Lemay et al., 2013), we find that arsenic perturbs lactose synthesis pathway via 4 genes, namely, GALK1, HK1, NME1-NME2 and SLC2A9 (Figure 5B; Supplementary Table S6). We have performed the same analysis for the chemicals compiled in ExHuMUS and ExHuM Explorer and their results are presented in Supplementary Table S6.

#### 3.7.2. Cytokine signalling pathway

Cytokines are small peptides which are involved in autocrine, paracrine, and endocrine signalling, and are responsible for the regulation of immune responses (Brooks et al., 2017). The presence of human milk contaminants influences the quality of the mother’s milk by causing a disruption in the cytokine production which can impact effective immune response in developing infants (Cohen, 1999; Rebelo and Caldas, 2016; Leibson et al., 2018; Pajewska-Szmyt et al., 2019). On analyzing the list of 116 cytokine receptors with chemical interactions obtained from ToxCast and CTD, we found that 22 chemicals compiled in ExHuMId interact with 32 cytokine receptors, which in turn could interfere with signalling or production of 64 cytokines (Methods; Figure 7; Supplementary Table S7). These interactions are displayed in the form of a tripartite network in Figure 7. Among the chemicals in ExHuMId, arsenic targets the highest number of cytokine receptors (24 genes) followed by Benzo[a]pyrene (9 genes). Among the cytokine receptors, CD40 is perturbed by 17 contaminants compiled in ExHuMId, and the binding of these contaminants to the CD40 receptor could interfere with the signalling and production of CD40LG, a cytokine specific to CD40 (Figure 7; Supplementary Table S7). Thus, human milk contaminants targeting cytokine receptors could bind to these receptors and interfere with normal function of cytokines. For the chemicals compiled in ExHuMUS and ExHuM Explorer, we have also performed the same analysis, and found several contaminants in these resources to be capable of influencing cytokine signalling and production (Supplementary Table S7).

#### 3.7.3. Influence on drug or xenobiotic transporters

The transport of contaminants into human milk can be both via diffusion or mediated by xenobiotic transporters (Ito and Alcorn, 2003; García-Lino et al., 2019). It has been reported that several membrane transporters belonging to the family of ABC and SLC transporters are expressed in mammary gland during lactation (García-Lino et al., 2019). In Alcorn *et al*. (Alcorn et al., 2002) study, 19 out of 30 transporters genes considered were found to be expressed in the mammary gland during lactation based on their Real-Time Reverse Transcription-Polymerase Chain Reaction (RT-PCR) analysis. The analysis of this dataset with chemical-gene interactions obtained from ToxCast and CTD revealed that 15 contaminants in ExHuMId target 9 transporters which are expressed during lactation (Methods; Figure 6B; Supplementary Table S8). Of these, there are two prominent transporter protein genes, namely, SLC22A1 and SLC22A4, which were found to be expressed 4-fold during lactation (Alcorn et al., 2002) (Figure 6B; Supplementary Table S8). Among the contaminants in ExHuMId, Arsenic targets 7 transporter genes. The ABCB1 transporter protein gene appears to be targeted by the maximum number of contaminants in ExHuMId (Figure 6B; Supplementary Table S8). We have also performed the same analysis for the chemicals compiled in ExHuMUS and ExHuM Explorer, and these results are presented in Supplementary Table S8.

From the analysis reported in this section, it is evident that the human milk contaminant Arsenic can target several genes or proteins in lactation pathway, cytokine signalling and production pathway, and xenobiotic transporters (Figures 5–7). Based on the compilation of studies in ExHuMId, Arsenic was detected in human milk samples collected from 3 states of India, namely, Chhattisgarh, Maharashtra and West Bengal. Arsenic was also found in a human milk sample from the United States, as reported in Lehmann *et al*. (Lehmann et al., 2018). From the evidence in scientific literature, Arsenic has been found to be present in many biospecimens and detected all around the world (Mandal, 2002). Especially, the primary source of Arsenic is known to be ground water or drinking water (https://www.who.int/news-room/fact-sheets/detail/arsenic) (Smith and Smith, 2004). Moreover, there are several studies which have reported on Arsenic contamination in ground water and drinking water samples collected from across several states of India (Bhattacharya et al., 1997; Borah et al., 2010; Sharma et al., 2013; Kumar et al., 2016). Thus, it is not surprising that Arsenic has been found as a human milk contaminant.

## 4. Conclusions

Human milk is the sole source of nourishment for infants for the first few months of their lives, during which exposure to environmental contaminants is a concern. These contaminants may have an impact on maternal health and lactation as well. Understanding the effects of these environmental contaminants to maternal and infant health remains challenging (Sonawane, 1995; Landrigan et al., 2002; Mead, 2008). In recent years there is an increased interest towards the development of an integrated approach in toxicology known as the exposome which captures all the environmental exposures of humans during their lifetime, and the implications of these exposures on their health (Vermeulen et al., 2020). In this work we have developed a comprehensive resource on <u>Ex</u>posome of <u>Hu</u>man <u>M</u>ilk across <u>I</u>n<u>d</u>ia, ExHuMId, through a systematic approach. ExHuMId compiles detailed information about 101 environmental contaminants detected in human milk samples studied across India from 36 published research articles. To build ExHuMId, we first performed an extensive PubMed literature mining which resulted in 1704 research articles relevant to human milk or breast milk from India. We then manually curated these 1704 research articles and identified 36 research articles containing the analytical studies specific to human milk contaminants in samples from India. Finally, from the curated list of 36 research articles, we extracted, compiled and unified information on the geographical location of the samples, maternal factors associated with the sample donors, and concentrations of contaminants detected in the human milk samples. ExHuMId compiles detailed information about contaminants detected in human milk samples across 13 states of India (Figure 2A) with a wide range of standardized maternal factors (Figure 2B). Furthermore, we have classified the 101 human milk contaminants based on their environmental sources (Figure 2C) and their chemical classification (Figure 2D). Lastly, we obtained 2D and 3D structure information for the contaminants, their physicochemical properties, predicted ADMET properties, molecular descriptors, and their target genes. ExHuMId contains all the above information for 101 environmental contaminants and is accessible online at: https://cb.imsc.res.in/exhumid/.

We have analyzed the human milk contaminants compiled in ExHuMId and two other resources from three perspectives. We first compared ExHuMId with the well-known chemical regulatory and guidelines lists to identify potential EDCs, carcinogens, neurotoxins or other hazardous chemicals. Of 101 human milk contaminants in ExHuMId, 43 were found to be potential EDCs, and 23 were found to be potential carcinogens. Among these 7 were found to be class 1 carcinogens, which according to the criteria provided by the IARC monographs on carcinogens (Samet et al., 2020), have sufficient evidence of carcinogenicity in humans. The 7 class 1 carcinogens detected in ExHuMId are: Arsenic, Benzo(a)pyrene, Cadmium, Lindane, Tetrachlorodibenzodioxin, 2,3,4,7,8-Pentachlorodibenzofuran and 3,4,5,3’,4’-Pentachlorobiphenyl. Lindane and Cadmium are also produced in high volume as per OECD and USHPV database (Methods). On analyzing the 101 human milk contaminants in ExHuMId, we found 14 potential neurotoxins detected from human milk samples collected across India, and of which 6 are produced in high volume. Similar analyses have been performed on the human milk contaminants compiled in ExHuMUS and ExHuM Explorer (Methods), and several chemicals of concern produced in high volume have been identified (Supplementary Table S3). Thus, from our systematic compilation and analysis of human milk contaminants, we observed there is a need for better chemical regulation and policy decisions to avoid these contaminants in human milk in India and globally.

The second perspective of our analysis enables to better understand the structural features and properties which influence the transfer of environmental contaminants into human milk, and thus, provides a way to predict the risk of contaminant entering human milk. Due to the lack of experimental data on M/P ratios of human milk contaminants in ExHuMId, we considered the dataset reported by Vasios *et al*. (Vasios et al., 2016) and performed a comparison of the physicochemical properties that have been widely reported to influence the transfer of contaminants or drugs into human milk. Through our analysis we observed that the distributions of physicochemical properties of contaminants in ExHuMId, ExHuMUS and ExHuM Explorer are close to the distributions of physicochemical properties of chemicals reported as highly likely to transfer to human milk in Vasios *et al*. (Vasios et al., 2016). This gives insights into the role of the structural features of contaminants that influence their transfer into human milk.

The third aspect of our analysis predicts the effect of the human milk contaminants on lactation pathway and cytokine signalling and production pathway, using a systems biology approach. Based on the interaction data obtained from ToxCast and CTD, we infer that many of the environmental contaminants compiled in the above-mentioned 3 datasets can interact with genes associated with lactation and cytokine signalling and production. Lastly, we analyzed the interactions obtained from ToxCast and CTD with transporter proteins reported to be expressed during lactation, and found several contaminants that show interactions; these contaminants may have a higher chances of transfer into human milk, due to their interactions with xenobiotic transporters. The above observations need to be critically validated using experimental approaches, which should encompass various disciplines, to understand the influence of environmental contaminants on maternal and infant health (Pajewska-Szmyt et al., 2019).

The development of a resource on human milk exposome specific to India is the first step in covering the wide range of information related to detected human milk contaminants, their concentrations, maternal factors, and other information which are dispersed across a large body of scientific literature. The determination of mean concentrations of contaminants or any established benchmarks like reference dose (RfD) or Tolerable Daily Intake (TDI) or Average Daily Dose (ADD) is not ventured into in this study, as the data compiled in this work is diverse in consonance with the breadth of the Indian population. It is important to highlight the availability of guidelines provided by the US Environmental Protection Agency (EPA) on Child-specific Exposure Scenarios Examples (U.S. EPA, 2014) for Indian scenarios, which can help to estimate the above benchmarks specific to India. During our literature mining we also found thousands of research articles available in the corpus of PubMed, on the detection of environmental contaminants in human milk across the world. Thus, the expansion of human milk exposome resources worldwide, and the availability of experimentally determined M/P ratio for environmental contaminants can help in better risk assessment and management of human milk contaminants. Importantly, further studies are necessary to understand the influence of variable factors such as maternal factors (Mead, 2008; Anadón et al., 2011; LaKind et al., 2018), the pharmacokinetics of environmental contaminants (Anadón et al., 2011; Lehmann et al., 2018), and the complexity of lactation pathways and physiology (Rebelo and Caldas, 2016; LaKind et al., 2018) in order to incorporate these variables in the risk estimation of human milk contaminants. We also note that there are several studies on detection of environmental contaminants in other specimens such as blood, plasma, serum, placenta, urine, saliva across India, and substantial manual effort is required to develop a comprehensive exposome resource specific to India which is beyond the scope of this work.

## CRediT author contribution statement

**Bagavathy Shanmugam Karthikeyan:** Conceptualization, Data curation, Formal analysis, Software, Writing – original draft. **Janani Ravichandran:** Conceptualization, Data curation, Formal analysis, Software, Visualization, Writing – original draft. **S. R. Aparna:** Formal analysis, Writing – original draft. **Areejit Samal:** Conceptualization, Investigation, Formal analysis, Supervision, Project administration, Funding acquisition, Writing – review & editing.

## Acknowledgements

AS would like to thank S. Krishnaswamy for discussion, and B. Raveendra Reddy for help in setting up the web server. AS acknowledges support from a Ramanujan fellowship (SB/S2/RJN-006/2014) from the Science and Engineering Research Board (SERB) India, the Department of Atomic Energy (DAE) India, and a Max Planck Partner Group from the Max Planck Society Germany. The funders have no role in study design, data collection, data analysis, manuscript preparation or decision to publish.

## Supplementary Tables

**Table S1:** List of 36 curated research articles containing the analytical studies that detected human milk contaminants from samples across India. This list of 36 curated research articles were filtered by manual curation of 1704 research articles obtained through literature mining.

**Table S2:** The list of 269 human milk contaminants in Global ExHuM which is the union of three resources, namely, ExHuMId, ExHuMUS, and ExHuM Explorer compiling 101, 127, and 183 contaminants, respectively. The subset of 44 contaminants that are present across all the three resources, namely, ExHuMId, ExHuMUS and ExHuM Explorer, is referred to as Common ExHuM.

**Table S3:** The table lists the 101 human milk contaminants from ExHuMId which are present across different categories of chemical lists including the lists of hazardous substances, substances specific to Indian regulations, and substances in possible everyday exposure.

**Table S4:** The table lists the 6 computed physicochemical properties for the chemicals compiled from four resources, namely, ExHuMId, ExHuMUS, ExHuM Explorer, and Vasios *et al*. (2016).

**Table S5:** The table lists the mean, standard deviation (SD), minimum value (Min), and maximum value (Max) for the 6 physicochemical properties, for the sets of human milk contaminants in ExHuMId, ExHuMUS, ExHuM Explorer, high risk (HR) compounds in Vasios *et al*. with M/P ratio ≥ 1, and low risk (LR) compounds in Vasios *et al*. with M/P ratio ≤ 0.25, ≤ 0.5, ≤ 0.75 and < 1.

**Table S6:** The table lists the genes or proteins involved in the lactation pathways which can be targets of human milk contaminants in Global ExHuM. The genes involved in lactation pathways namely, prolactin and oxytocin signalling pathways, and lactose synthesis pathway were compiled from KEGG and NetPath, while the contaminant-gene interactions were compiled from CTD and ToxCast.

**Table S7:** The table gives the tripartite network of human milk contaminants in Global ExHuM, cytokine receptors, and cytokines. The cytokine receptors were compiled from Book NBK6294, Guide to Pharmacology, HGNC, and KEGG, while the contaminant-cytokine receptor interactions were compiled from CTD and ToxCast.

**Table S8:** The table lists the genes involved in the xenobiotic transport which can be targets of human milk contaminants in Global ExHuM. The genes involved in xenobiotic transport were compiled from Alcorn *et al*. (2002), while the contaminant-gene interactions were compiled from CTD and ToxCast.

